# *SRP19* and the protein secretion machinery is a targetable vulnerability in cancers with *APC* loss

**DOI:** 10.1101/2024.01.30.578083

**Authors:** Xinqi Xi, Ling Liu, Natasha Tuano, Julien Tailhades, Dmitri Mouradov, Oliver Sieber, Max Cryle, Eva Segelov, Joseph Rosenbluh

## Abstract

Adenomatous Polyposis Coli (*APC*) is a tumour suppressor that is frequently lost in colorectal and other cancers. A common mechanism for *APC* loss includes heterozygous *APC* deletion. Here, we show that *SRP19*, is located near *APC* and is often co-deleted in these tumours. Heterozygous *APC/SRP19* loss leads to lower levels of *SRP19* mRNA and protein. Consequently, cells with *APC/SRP19* loss are vulnerable to partial suppression of *SRP19*. We show that *SRP19* loss is a unique vulnerability since *SRP19* is rate limiting for the formation of the Signal Recognition Particle (SRP), a complex that mediates translocation of proteins to the ER. Consistent with these observations, partial *SRP19* knock-down or low dose Arsenic Trioxide treatment induces ER stress and inhibits proliferation in *APC/SRP19* loss cancers. Our work identifies a new strategy to treat cancers with *APC/SRP19* heterozygous deletions and provides a framework for identifying vulnerabilities associated with loss of a tumour suppressor.

## Introduction

*APC* is part of the destruction complex that mediates β-catenin phosphorylation leading to its degradation(1). Loss of function genomic alterations in *APC* are frequent in colorectal cancer(2,3) but, are also found in other cancers such as gastric(4) and breast(5) tumours. Two main mechanisms contribute to *APC* loss in cancer, mutations and/or deletions. *APC* loss of function is found in ∼80% of colorectal cancers, with ∼20% of these cases involving heterozygous *APC* gene deletions and mutation of the other allele(6). Studies in animal models and colon organoids demonstrated that even in advanced carcinomas restoration of *APC* leads to cell differentiation and tumour regression(7), demonstrating that supressing *APC* activity is always required in these tumours for tumour maintenance. However, restoration of *APC* activity is challenging to achieve in a therapeutic setting, and we currently lack strategies for direct targeting tumour suppressor genes (TSGs), such as *APC,* that are lost in cancer.

Although DNA copy number loss cannot be directly targeted, previous studies have shown that heterozygous deletion of a genomic region including a TSG can result in collateral loss of nearby essential genes that become vulnerabilities in these cancers(8-11). This phenomenon known as CYCLOPS(8) has been shown in several contexts. For example, heterozygous deletion of *PSMC2,* a cell essential proteasomal gene, is frequent in cancer leading to a unique vulnerability to partial inhibition of *PSMC2*(8). Paolella et al. showed that the cell essential splicing factor, *SF3B1* is a dependency in cancers with loss of *SF3B1*(12). Liu et al. demonstrated that *TP53* deletion results in heterozygous loss of the cell essential transcription regulator *POLR2A* and that POLR2A inhibitors are effective in treating these cancers(10). Another example of a druggable CYCLOPS is *CSKN1A1* the target of Lenalidomide in myelodysplastic syndrome (MDS) with deletion of chromosome 5q(13).

SRP19, is a component of the Signal Recognition Particle (SRP), an evolutionary conserved cellular complex that mediates transport of proteins to the endoplasmic reticulum (ER)(14). The SRP complex is a ribonucleoprotein complex composed of a non-coding RNA (7SL) and six proteins. SRP19 interacts with the 7SL RNA and forms a complex that can recognize a signal sequence on the emerging protein chain. When a protein signal is recognized, the SRP complex binds to the signal peptide through SRP54, temporarily halting translation and facilitating the translocation of the ribosome-nascent chain complex to the ER.

Here, we show that *SRP19* is frequently co-deleted in the cancers with heterozygous loss of *APC*. We show that *SRP19* is a central component of the SRP complex and that loss of *SRP19* results in destabilisation of the SRP complex. We demonstrate that the protein secretion machinery is a unique targetable vulnerability in cancers harbouring *APC/SRP19* loss.

## Results

### *SRP19* mRNA and protein levels are closely associated with *APC* loss in cultured cell lines and patient tumours

To identify gene expression changes that are associated with *APC* copy number loss, we used the Cancer Cell Line Encyclopedia (CCLE) database(15). We calculated a Pearson correlation between *APC* copy number variation (CNV) and gene expression for every gene in the human genome across 1,440 cell lines (Fig. 1A and Supplementary Table S1). We found that expression of genes located on chromosome 5 near *APC* (e.g. *SRP19*, *ATG12*, *PHAX*, *REEP5*, *TMED7* and *COMMD10*) were the most highly correlated with *APC* CNV (Fig. 1A), suggesting that the expression of these genes changes is regulated by their CNV. Among these genes, *SRP19* is the closest to *APC* (15kb apart) and *SRP19* expression had the highest correlation with *APC* CNV in cell lines (Fig. 1A,B). These results are consistent with previous reports showing that CNV and mRNA expression are highly correlated for ∼50% of genes in the human genome(8).

**Figure 1:**
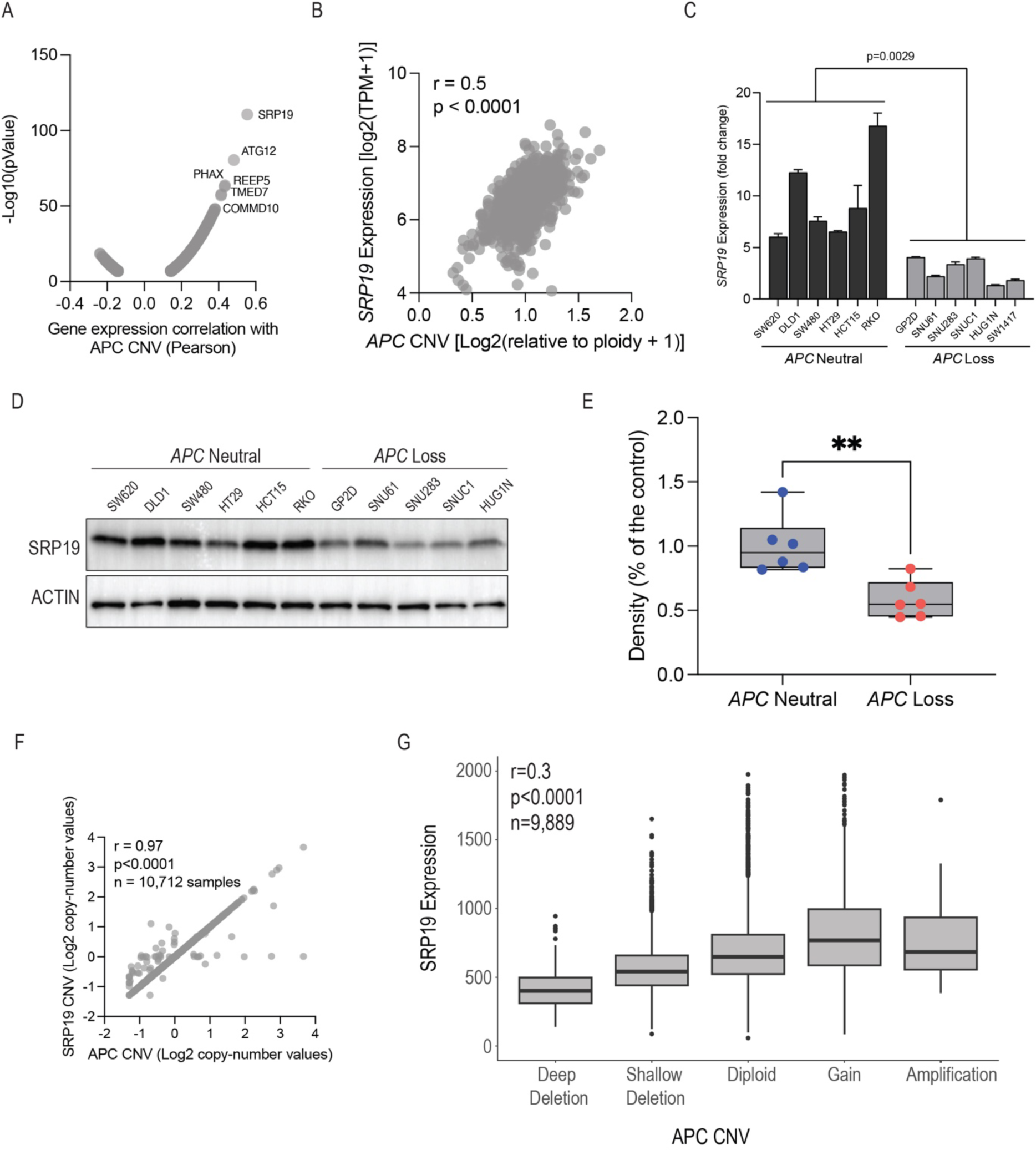
*APC* loss leads to loss of *SRP19* DNA, mRNA and protein. (A) Volcano plot showing genes whose mRNA expression is correlated with *APC* copy number. Indicated genes are located on chromosome 5. (B) Correlation between *SRP19* expression and *APC* copy number. Each dot represents a cancer cell line. (C) qRT-PCR measuring SRP19 mRNA in cells with neutral *APC* copy number or cell-lines with heterozygous *APC* copy number loss. p.Value was calculated using a two-tailed unpaired T.test. (D) SRP19 protein levels in cells with neutral or heterozygous loss of *APC* copy number. (E) Quantification of SRP19 protein levels from three biological replicates (Supplementary Fig. 1). p.Value was calculated using a two-tailed unpaired T.test. (F) Correlation between *SRP19* and *APC* copy number across 10,712 samples from TCGA. (G) *SRP19* mRNA levels in 9,889 patient samples from TCGA with various levels of *APC* copy number. Definitions of *APC* CNV are from the cBio portal(17) and are based on the GISTIC copy number values (Deep deletion -2, Shallow deletion -1, Diploid 0, Gain 1, Amplification 2). Correlation and p.Value calculated using Pearson correlation.

To validate these observations we quantified the levels of *SRP19* mRNA (Fig. 1C) and protein (Fig. 1D,E and Supplementary Fig. 1) in a panel of cancer cell lines and found that cell lines with heterozygous *APC* loss have lower levels of *SRP19* mRNA and protein. To confirm these observations in patient tumours we analysed The Cancer Genome Atlas (TCGA) dataset(15,16). We found that across 10,712 cancer samples, heterozygous loss of *APC* was almost always accompanied by heterozygous *SRP19* loss (r=0.97, p<0.0001, Fig. 1F). Furthermore, we found that across 9,889 tumours, that have CNV and gene expression data available, *SRP19* expression was highly correlated with *APC* CNV (r=0.3, p<0.0001, Fig. 1G). Based on these observations we conclude that heterozygous loss of *APC* leads to heterozygous loss of *SRP19* and to decreased levels of *SRP19* mRNA and protein in both cell lines and patient tumours.

### Cells harbouring *APC* copy number loss are highly sensitive to partial suppression of *SRP19* expression

*SRP19* is a core cell-essential gene that is required for proliferation in all cell types in a context independent manner(9). CRISPR mediated knockout of *SRP19* across 1,019 cell lines, results in cell death in all cell lines (Supplementary Fig. 2A) that is not correlated to *SRP19* expression levels (Supplementary Fig. 2B). CRISPR knockout experiments result in complete loss of protein expression(18) and are not able to identify context specific vulnerabilities that are associated with partial loss of gene expression(9). Since *SRP19* is a core essential gene, we hypothesised that cancers with low levels of *SRP19* mRNA and protein will be more sensitive to partial inhibition of *SRP19* expression. Using the DepMap RNAi dataset(19), we found that shRNA mediated suppression of *SRP19* across a panel of 657 cell lines was positively correlated with *SRP19* gene expression (r=0.2, p<0.0001, Supplementary Fig. 2C) suggesting partial *SRP19* knockdown is a vulnerability in cells with loss of *APC/SRP19*.

Since it is difficult to predict or titre the level of gene suppression with shRNAs, we used siRNAs to validate these observations. We designed two *SRP19* targeting siRNAs that reduce but do not eliminate *SRP19* expression (Fig. 2A). To assess partial *SRP19* as a target we selected a panel of cancer cell lines harbouring heterozygous loss or neutral copy number of *APC/SRP19*. Following transfection with *SRP19* targeting siRNAs cells were incubated for 7 days and proliferation was measured using crystal violet staining (Fig. 2B and Supplementary Fig. 2D). We found that while cancer cells harbouring *APC/SRP19* loss were highly sensitive to partial suppression of *SRP19,* cells with neutral *APC/SRP19* copy number could tolerate partial suppression of *SRP19* (Fig. 2C), suggesting partial suppression of *SRP19* expression as a vulnerability in *APC/SRP19* loss cancers. RNAi off-target effects are a major hurdle in interpretation of RNAi experiments(19). To ensure siRNA specificity we used a phenotypic rescue experiment. Following overexpression of *SRP19* in SNU61 or SW1463, two *APC/SRP19* loss colorectal cancer cell lines, cells were transfected with *SRP19* targeting siRNAs and after 7 days we measured SRP19 protein levels and cell proliferation. Overexpression of *SRP19* inhibited siRNA mediated suppression of SRP19 protein levels (Fig. 2D and Supplementary Fig. 2E) and its effect on proliferation (Fig. 2E and Supplementary Fig. 2F-H). These results demonstrate the specificity of *SRP19* targeting siRNAs and identify *SRP19* as a dose dependent vulnerability in cell lines harbouring *APC/SRP19* loss.

**Figure 2:**
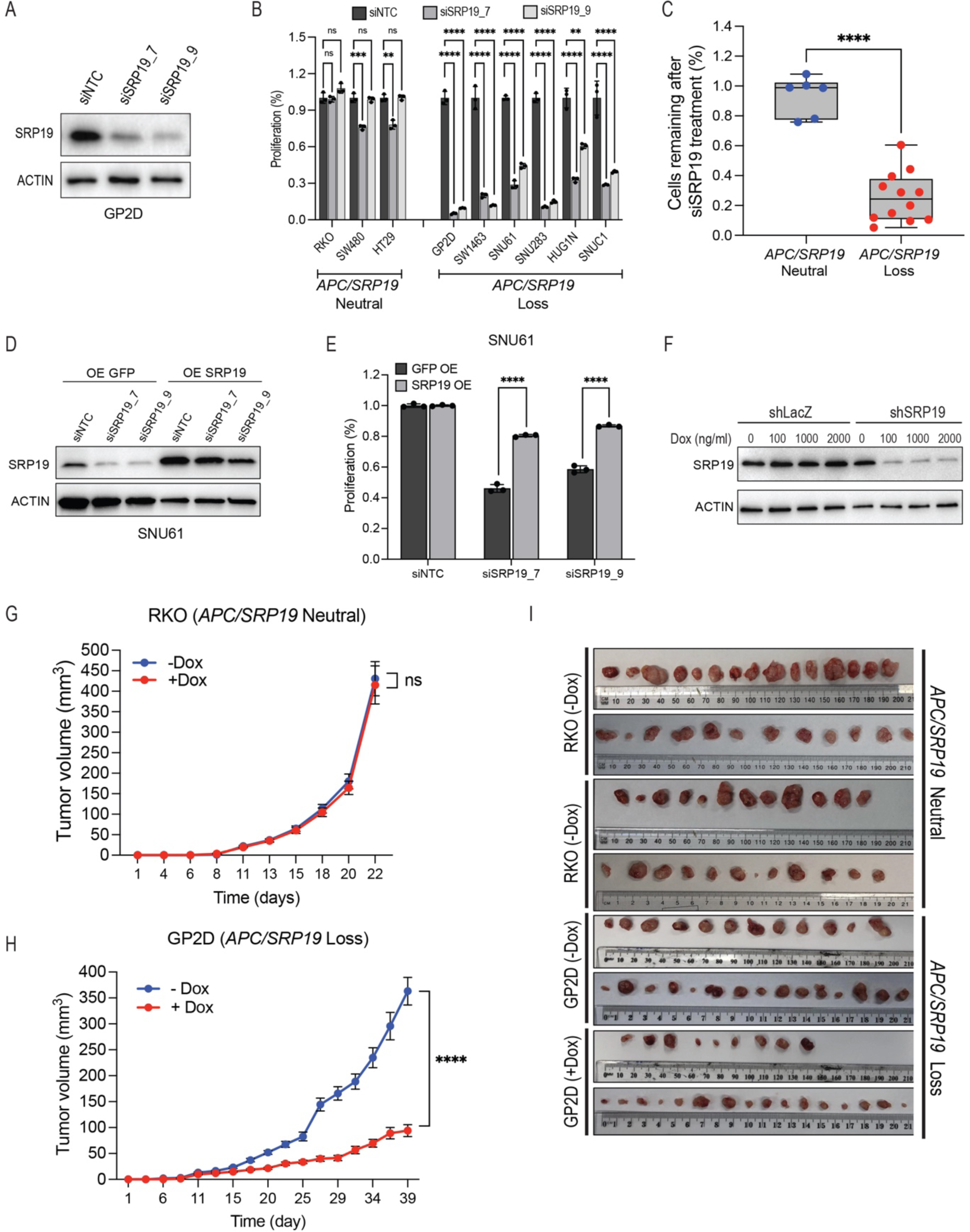
Cells with *SRP19* loss are highly sensitive to further suppression of *SRP19* expression. (A) SRP19 protein levels 3 days post transfection with *SRP19* targeting siRNAs in GP2D cells. (B) Cell proliferation measured using crystal violet staining 7 days post transfection with *SRP19* targeting siRNAs in cell lines with neutral or loss of *APC/SRP19* copy number. pValue calculated using a two-tailed unpaired T.test. (C) Comparison of cell viability following suppression of *SRP19* expression in cells with neutral or loss of *APC/SRP19* copy number. pValue calculated using a two-tailed unpaired T.test. (D) SRP19 protein levels in SNU61 (*APC/SRP19* loss) following overexpression of *SRP19* and transfection of *SRP19* targeting siRNAs. (E) Proliferation following rescue of *SRP19* expression measured by crystal violet staining in SNU61 cells. pValue calculated using a two-tailed unpaired T.test. (F) SRP19 protein levels in GP2D cells containing inducible *SRP19* shRNA following Doxycycline treatment. (G) Tumour growth in mouse xenografts injected with RKO cells containing an inducible *SRP19* targeting shRNA treated with or without Dox. pValue calculated using a two-tailed unpaired T.test. (H) Tumour growth in mouse xenografts injected with GP2D cells containing an inducible *SRP19* targeting shRNA treated with or without Dox. pValue calculated using a two-tailed unpaired T.test. (I) Images of tumours from (G) and (H).

To assess partial *SRP19* knockdown as a vulnerability, associated with *APC/SRP19* loss, in an in vivo setting, we used a mouse xenograft model and a doxycycline (DOX) inducible shRNA expression system(20) for partial suppression of *SRP19* expression (Fig. 2F). Immuno-deficient NOD-SCID mice were injected with GP2D (heterozygous *APC/SRP19* loss colon cancer) or RKO (*APC/SRP19* neutral colon cancer) cells containing a DOX inducible *SRP19* targeting shRNA. Two weeks after injection mice developed a palatable tumour and were than given normal or DOX containing food. DOX treated tumours, extracted from animals, had reduced levels of SRP19 protein (Supplementary Fig. 2I). We found that in RKO tumours were not affected by partial suppression of *SRP19* expression (Fig. 2G,I and Supplementary Fig. 2J) and GP2D tumours were highly sensitive to RNAi mediated partial suppression of *SRP19* expression (Fig. 2H,I and Supplementary Fig. 2K). Based on these observations, we conclude that partial suppression of *SRP19* is a unique vulnerability in *APC/SRP19* loss cancers.

### *SRP54* and *SRP68* are vulnerabilities in *APC/SRP19* loss cancers

SRP19 is part of the SRP complex which is a core cell essential complex that plays a central role in translocation of newly translated proteins to the ER ((14) and Fig. 3A). Confirming that the SRP complex is a core cell essential complex, CRISPR knockout experiments (DepMap) show that all six SRP complex genes as core cell-essential genes that inhibit proliferation in a context independent manner (Supplementary Fig. 3A). Fitness screens are effective in identifying gene-gene relationships and uncovering complex pathway networks(21). Recent studies show that genome wide RNAi loss of function proliferation screens induce partial gene knockdown and can identify correlated pathway dependencies for core cell essential genes(18). To identify gene dependencies that correlate with *SRP19* dependency, for each gene in the human genome, we calculated a Pearson correlation with *SRP19* dependency, using the DepMap shRNA dataset across 657 cancer cell lines (Fig. 3B and Supplementary Table S2). To identify enriched pathways, we selected genes with a Pearson correlation >0.2 or <-0.2 and used MsigDB(22) pathway enrichment analysis. We found that the SRP complex signature was positively correlated with *SRP19* dependency, and that components of the ribosome and translation machinery were negatively correlated with *SRP19* dependency (Fig. 3B,C). Since the SRP complex directly binds to the ribosome(23-25), it is likely that cells with low *SRP19* have more ribosomes available and are thus less sensitive to partial inhibition of ribosomal subunits.

**Figure 3:**
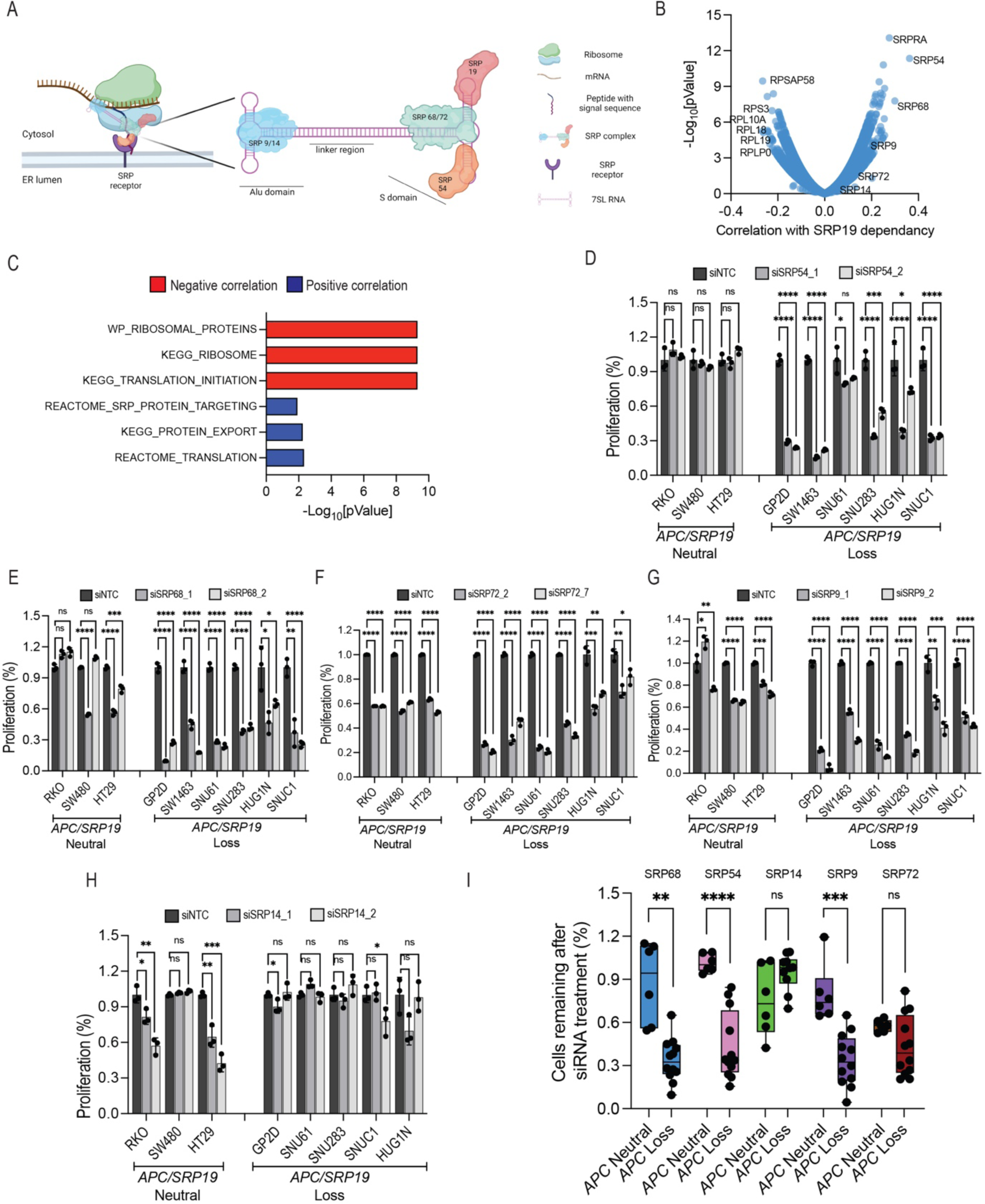
*SRP68* and *SRP54* are also vulnerabilities in SRP19 loss cancers. (A) Illustration of the SRP complex. (B) Volcano plot showing gene dependencies that correlate or anti-correlate with SRP19 RNAi dependency across the DepMAp dataset. (C) Geneset enrichment analysis of dependencies that are positively (blue) or negatively (red) correlated with *SRP19* dependency. (D) Cell proliferation measured using crystal violet staining 7 days post infection with *SRP54* targeting siRNAs in cell lines with neutral or loss of *APC/SRP19*. pValue calculated using a two-tailed unpaired T.test. (E) Cell proliferation measured using crystal violet staining 7 days post infection with *SRP68* targeting siRNAs in cell lines with neutral or loss of *APC*. pValue calculated using a two-tailed unpaired T.test. (F) Cell proliferation measured using crystal violet staining 7 days post infection with *SRP72* targeting siRNAs in cell lines with neutral or loss of *APC*. pValue calculated using a two-tailed unpaired T.test. (G) Cell proliferation measured using crystal violet staining 7 days post infection with *SRP9* targeting siRNAs in cell lines with neutral or loss of *APC*. pValue calculated using a two-tailed unpaired T.test. (H) Cell proliferation measured using crystal violet staining 7 days post infection with *SRP14* targeting siRNAs in cell lines with neutral or loss of *APC*. pValue calculated using a two-tailed unpaired T.test. (I) Comparison of SRP complex dependencies between cell lines with neutral or loss of *APC/SRP19*. pValue calculated using a two-tailed unpaired T.test.

SRP complex proteins that are part of the S domain (Fig. 3A) were more correlated with *SRP19* dependency than proteins that are part of the Alu domain (Fig. 3B and Supplementary Fig. 3B-F and Supplementary Table S2). Since SRP19 is also part of the S domain these results indicate that SRP19 is important for stability of S domain proteins. To assess if partial suppression of other SRP complex proteins is a vulnerability in cells with *APC/SRP19* loss, we designed siRNAs for all the components of the SRP complex (Supplementary Fig. 3G-K and Fig. 2A). Using a panel of cancer cell lines with neutral or loss of *APC/SRP19* copy number we assessed the effect of partial suppression of these genes on proliferation (Fig. 3D-I). Consistent with DepMap analysis (Fig. 3B), we found that siRNA mediated partial suppression of *SRP54* (Fig. 3D and Supplementary Fig. 3L) and *SRP68* (Fig. 3E and Supplementary Fig. 3M) inhibited proliferation only in *APC/SRP19* loss cells (Fig. 3I). Partial suppression of the Alu domain protein *SRP9* was also more dependent in cells with *APC/SRP19* loss (Fig. 3G,I and Supplementary Fig. 3O) and partial suppression of *SRP14* or *SRP72* did not show any differential sensitivity (Fig. 3F,H,I and Supplementary Fig. 3N,P). These results are consistent with structural studies showing that SRP19 binding to the 7SL RNA results in a structural change that is critical for binding of SRP54 and SRP68(26). Based on these findings we suggest a model in which SRP19 binding to the 7SL RNA is critical for stabilisation of the 7SL RNA S domain and recruitment of SRP68 and SRP54 (Fig. 3A). Cancers harbouring *APC/SRP19* loss have low levels of *SRP19* and as a result are sensitive to loss of *SRP68* and *SRP54*.

### SRP54 and SRP68 protein stability is regulated by SRP19 and altered in *APC/SRP19* loss cells

Our results demonstrate that partial suppression of *SRP54* and *SRP68* is a vulnerability in cells harbouring *APC/SRP19* copy number loss. Since all SRP complex genes are located on different chromosomes (*SRP19* (chr5), *SRP54* (chr14), *SRP68* (chr17), *SRP72* (chr4), *SRP14* (chr15), *SRP9* (chr1)) it is not likely that *SRP54* and *SRP68* dependency in cells with *APC/SRP19* loss is related to changes in their DNA copy number. To gain insights into why *SRP54* and *SRP68* are dependencies in cells with *APC/SRP19* loss, we measured mRNA and protein levels of SRP complex genes that scored as vulnerabilities in cells with *APC/SRP19* copy number loss. Using the CCLE dataset across 1,440 cell lines we found that in contrast to *SRP19* (Fig. 1B), there was no gene expression correlation between *APC/SRP19* copy number and gene expression of these SRP complex genes (Supplementary Fig. 4A-E). We further validated these observations using qRT-PCR on a panel of 8 cancer cell lines (Supplementary Fig. 4F), demonstrating that *SRP54* and *SRP68* dependency in *APC/SRP19* loss cells is not due to transcription regulation.

Different to mRNA, we found that SRP54 protein levels were reduced in cells with *APC/SRP19* loss (Fig. 4A). These observations suggest a model were SRP54 binding to the 7SL RNA is dependent on SRP19(26). In cells with low *SRP19* the free SRP54 that is not bound to the 7SL RNA and is degraded, resulting in low SRP54 protein levels and a dependency on partial *SRP54* knockdown. To directly assess this model, we measured the abundance of SRP complex proteins following partial suppression of *SRP19* (Fig. 4B). Consistent with our model, suppression of *SRP19* led to reduction in levels of *SRP54* protein in both *APC/SPR19* loss and neutral cells (Fig. 4B). In contrast, partial suppression of *SRP54* did not have any effect on the levels of SRP19 protein or other component of the SRP complex (Supplementary Fig. 4G). These results demonstrate that as predicted from structural models(26) following SRP19 binding to the 7SL RNA a structural change is induced leading to SRP54 binding. Thus, SRP54 stability is dependent on SRP19 protein levels and access unbound SRP54 is degraded in cells with *APC/SRP19* loss. To validate this model, we measured SRP54 protein levels in HT29 (*APC/SRP19* neutral) or GP2D (*APC/SRP19* loss) following treatment with Bortezomib, a proteosome inhibitor. We found that inhibiting the proteosome stabilised SRP54 protein levels only in an *APC/SRP19* loss cell demonstrating that SRP54 protein not bound to the 7SL complex in *APC/SRP19* loss cells is degraded by the proteosome (Fig. 4C).

**Figure 4:**
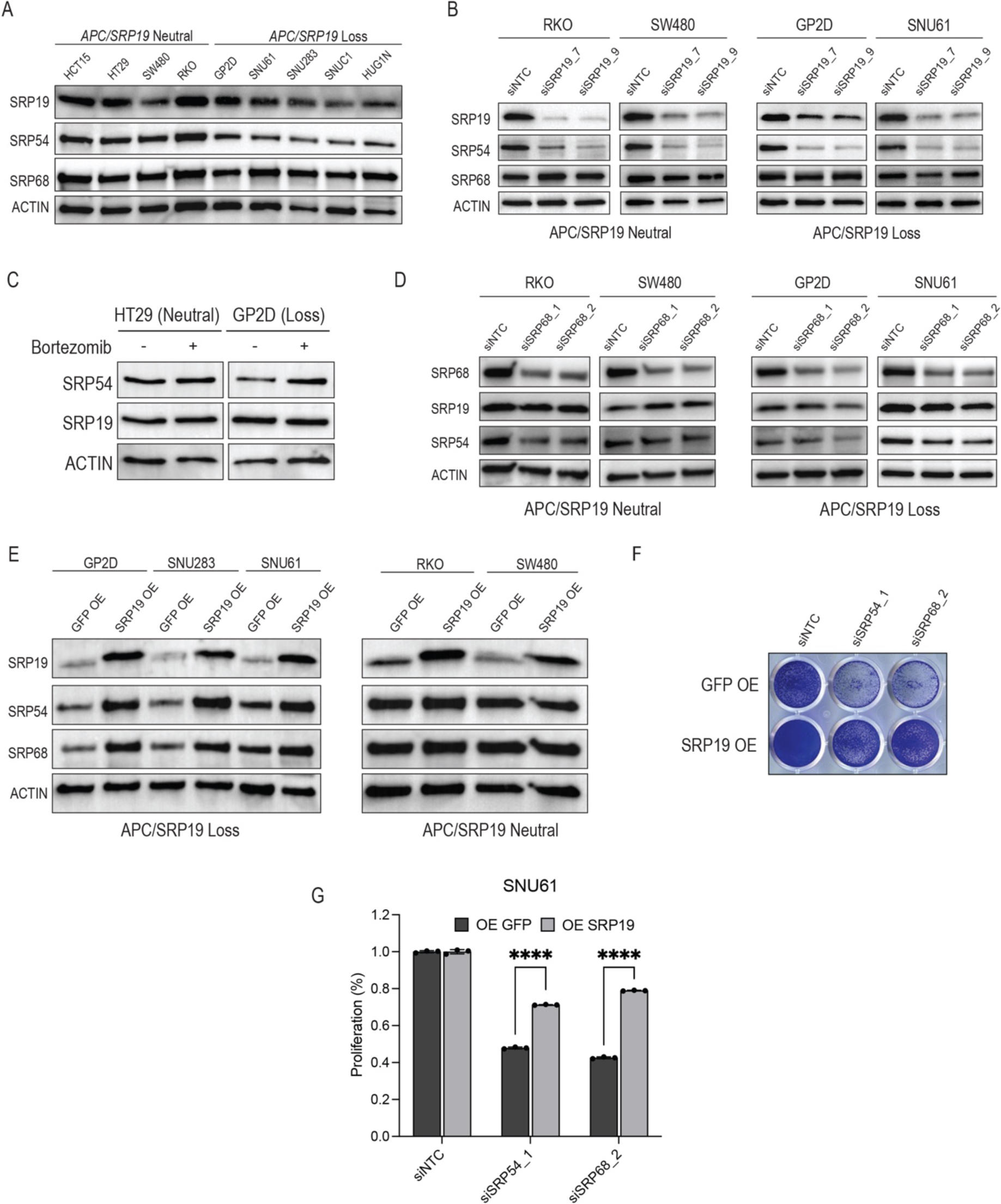
Loss of *APC/SRP19* destabilises components of the SRP complex. **(A)** Protein levels of SRP complex components in cells with *APC/SRP19* copy number neutral or loss. (B) Protein levels of SRP54 and SRP68 3 days post transfection with *SRP19* targeting siRNAs in *APC/SRP19* neutral or loss cells. (C) Protein levels of SRP19 and SRP54 in HT29 and RKO cells after 8h treatment with 100nM of bortezomib. (D) Protein levels of SRP19 and SRP54 3 days post transfection with *SRP68* targeting siRNAs in *APC/SRP19* neutral or loss cells. (E) SRP54 and SRP68 protein levels following overexpression of *SRP19* in *APC/SRP19* loss cells. (F) Crystal violet images of SNU61 cells overexpressing SRP19 or GFP 7 days post transfection with control or *SRP19* or *SRP54* targeting siRNA. (G) Quantification of crystal violet images in (F). pValue calculated using a two-tailed unpaired T.test.

*SRP68* also scored as an *SRP19* co-dependency (Fig. 3B,E). Structural studies have shown that SRP68 protein binds to the 7SL RNA and interacts with SRP54 and SRP72. SRP68 mRNA (Supplementary Fig. 4B) and protein (Fig. 4A) were not changed in *APC/SRP19* loss cells. To identify mechanisms associated with *SRP68* dependency we measured levels of SRP complex components following partial suppression of *SRP68* (Fig. 4D). We found that in both *APC/SRP19* loss and neutral cells partial suppression of *SRP68* reduced levels of SRP54 protein (Fig. 4D). These observations suggest that SRP68 requires interactions with SRP54 to assemble an active SRP complex. Due to the low levels of SRP54 found in cells with *APC/SRP19* loss, only a small portion of SRP68 is in an active complex resulting in increased sensitivity to suppression of *SRP68*.

To further validate that SRP54 and SRP68 protein stability is regulated by SRP19 and that low levels of SRP19 protein are important for regulation of SRP54 and SRP68 protein levels we used a rescue experiment. Following *SRP19* overexpression in cells with *APC/SRP19* loss or neutral copy number we measured levels of SRP54 and SRP68 proteins. We found that in consonance with our model SRP54 and SRP68 protein levels were increased following overexpression of *SRP19* only in cells with *APC/SRP19* loss (Fig. 4E). Furthermore, *SRP19* overexpression rescued the proliferation effect of siRNA mediated suppression of *SRP54* and *SRP68* in SNU61, (Fig. 4F,G) or GP2D (Supplementary Fig. 4H), two *APC/SRP19* loss colon cancer cell lines. Based on these observations we conclude that SRP19 protein levels regulate the stability and activity of SRP54 and SRP68. These results suggest that *APC/SRP19* loss cells will have less active SRP complex activity and thus less ability to secrete proteins.

### *APC/SRP19* loss cells have lower SRP complex activity and less protein secretion

To assess the activity of the SRP complex in *APC/SRP19* loss and neutral cells, we modified a previously described assay using Secreted Embryonic Alkaline Phosphatase (SEAP)(27). In this assay, a modified secreted version of the alkaline phosphatase SEAP is transported to the ER through the SRP complex and secreted to the media(28). To enable efficient monitoring of the protein secretion machinery we generated a lentiviral delivered secreted SEAP protein. Following transduction and Puromycin selection, an alkaline phosphatase luminesces assay is used to measure secreted SEAP in the media (Fig. 5A). To validate the ability of this assay to measure SRP complex activity we monitored SEAP secretion following siRNA mediated knockdown of SRP complex genes (Fig. 5B). As a positive control we used Brefeldin A (BFA), a small molecule toxin produced by fungi, that inhibits trafficking from the ER to the golgi(29). Consistent with previous reports(28), BFA treatment or suppression of *SRP19* or *SRP54* inhibited SEAP secretion (Fig. 5B), demonstrating the ability of this assay to monitor SRP complex activity and protein secretion.

**Figure 5:**
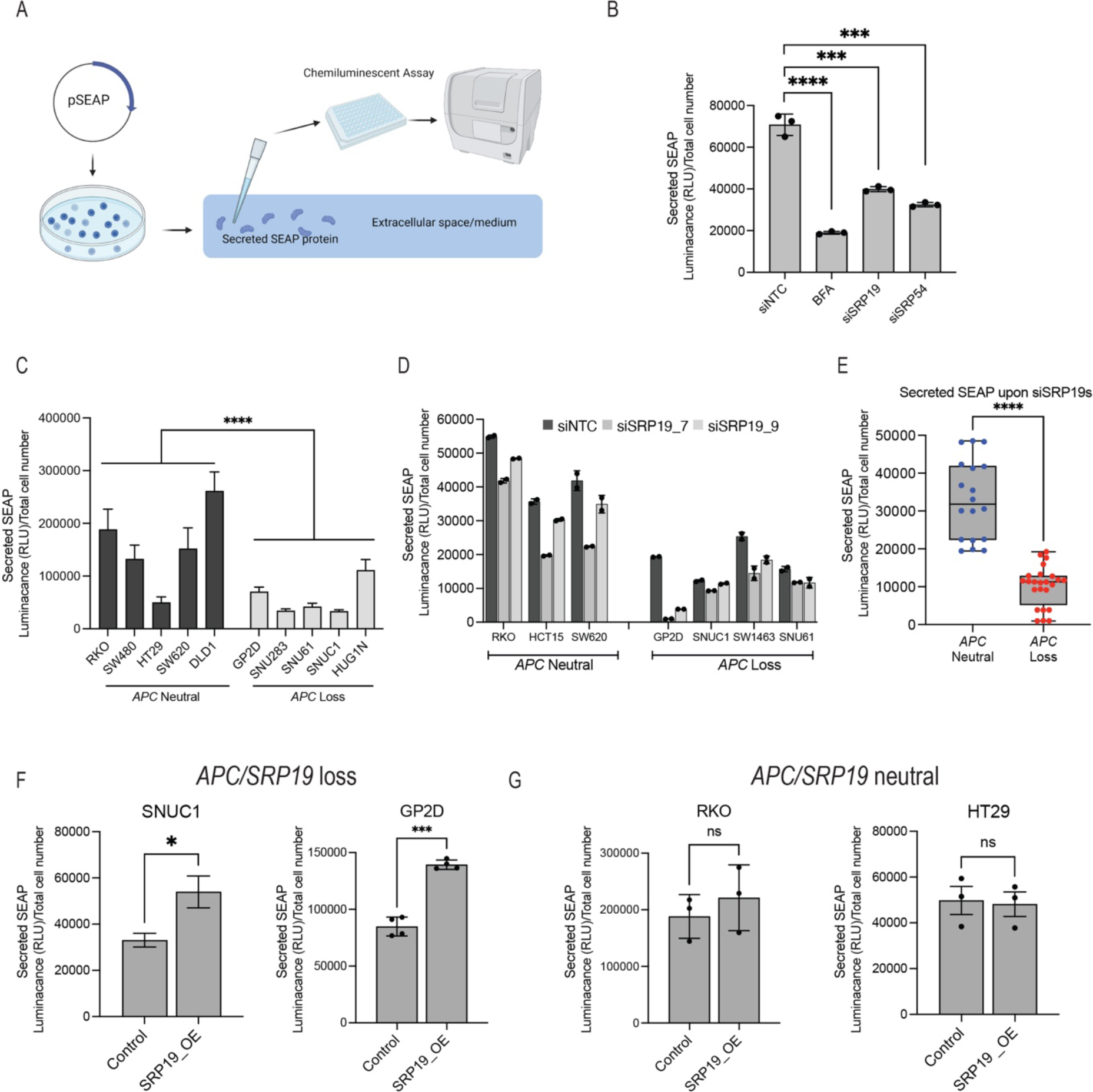
Cells harbouring *APC/SRP19* loss have lower levels of protein secretion activity. (A) Illustration of SEAP secretion assay system used to measure protein secretion. (B) SEAP secretion activity in GP2D cells transfected with the indicated siRNAs or treated for 2h with 7µM of BFA. p.Value calculated using a two-tailed unpaired T.test. (C) Basal SEAP protein secretion activity in cell lines with neutral or loss of *APC/SRP19* copy number stably expressing the SEAP reporter. SEAP activity was normalised to cell number. p.Value calculated using a two-tailed unpaired T.test. (D) SEAP activity measured in cells with neutral or loss of *APC/SRP19* following transfection with control or *SRP19* targeting siRNAs. p.Value calculated using a two-tailed unpaired T.test. (E) Comparison of secreted SEAP activity following partial suppression of *SRP19* in cells with neutral or loss of *APC/SRP19* copy number. p.Value calculated using a two-tailed unpaired T.test. (F) Rescue of SEAP secretion activity following overexpression of *SRP19* in cells with *APC/SRP19* copy number loss. p.Value calculated using a two-tailed unpaired T.test. (G) Rescue of SEAP secretion activity following overexpression of *SRP19* in cells with neutral *APC/SRP19* copy number. p.Value calculated using a two-tailed unpaired T.test.

To compare protein secretion activity between *APC/SRP19* neutral and loss cells we transduced 10 cell lines (5-neutral and 5-loss of *APC/SRP19* copy number) with the SEAP reporter and measured SEAP secretion. We found that *APC/SRP19* loss cells had lower basal protein secretion activity (Fig. 5C), suggesting that the low levels of *SRP19* and SRP complex in these cells leads to lower activity of the protein secretion machinery. To directly assess the effect of *SRP19* partial loss on the activity of the SRP complex we measured SEAP secretion after treatment with *SRP19* targeting siRNAs. We found that partial suppression of *SRP19* had a small effect on SEAP secretion in *APC* neutral cells (Fig. 5D) and was significantly inhibited in *APC/SRP19* loss cells (Fig. 5D,E). These observations are consistent with our previous findings suggesting that *SRP19* level is a key factor in regulating the SRP complex activity. To directly assess if lower levels of *SRP19* drive low protein secretion activity, we used a rescue experiment. Following overexpression of *SRP19* in *SRP19* loss cells we observed a ∼2 fold increase in SEAP secretion (Fig. 5F). SRP19 overexpression had no effect on SEAP secretion in *APC/SRP19* neutral cell lines (Fig. 5G) demonstrating that SRP19 is rate limiting for protein secretion and that the lower protein secretion activity in these cells is a consequence of the lower SRP19 protein levels and lower SRP complex activity in cells with *APC/SRP19* copy number loss.

### Induction of ER stress by Arsenic Trioxide is a vulnerability in *APC/SRP19* loss cancers

Our results demonstrate that cells with heterozygous loss of *APC/SRP19* have lower protein secretion activity due to lower levels of the SRP complex activity. These results suggest that *APC/SRP19* loss cells will have higher levels of ER stress and will be more sensitive to small molecules that induce ER stress or inhibit protein secretion. BiP (also known as GRP78 or HSPA5) is a protein chaperon that is upregulated during ER stress(30). Using a panel of cell-lines we found that basal levels of BiP were higher in most *APC/SRP19* loss cells compare to cells with neutral *APC/SRP19* copy number (Fig. 6A), suggesting that cells with *SRP19* loss have higher basal levels of ER stress. Consistent with these observations we found that following treatment with BFA, a known inducer of ER stress(28), BiP protein levels were up regulated more rapidly in *APC/SRP19* loss cells demonstrating that these cells are more sensitive to induction of ER stress (Fig. 6B). Furthermore, we found that siRNA mediated suppression of *SRP19* had a transient effect on BiP levels in a cell with neutral *APC/SRP19* copy number and that high BiP levels were maintained for longer periods in *APC/SRP19* loss cells (Fig. 6C). These results demonstrate that *SRP19* loss leads to increased basal levels of ER stress and suggest ER stress inhibitors as a vulnerability in these cells.

**Figure 6:**
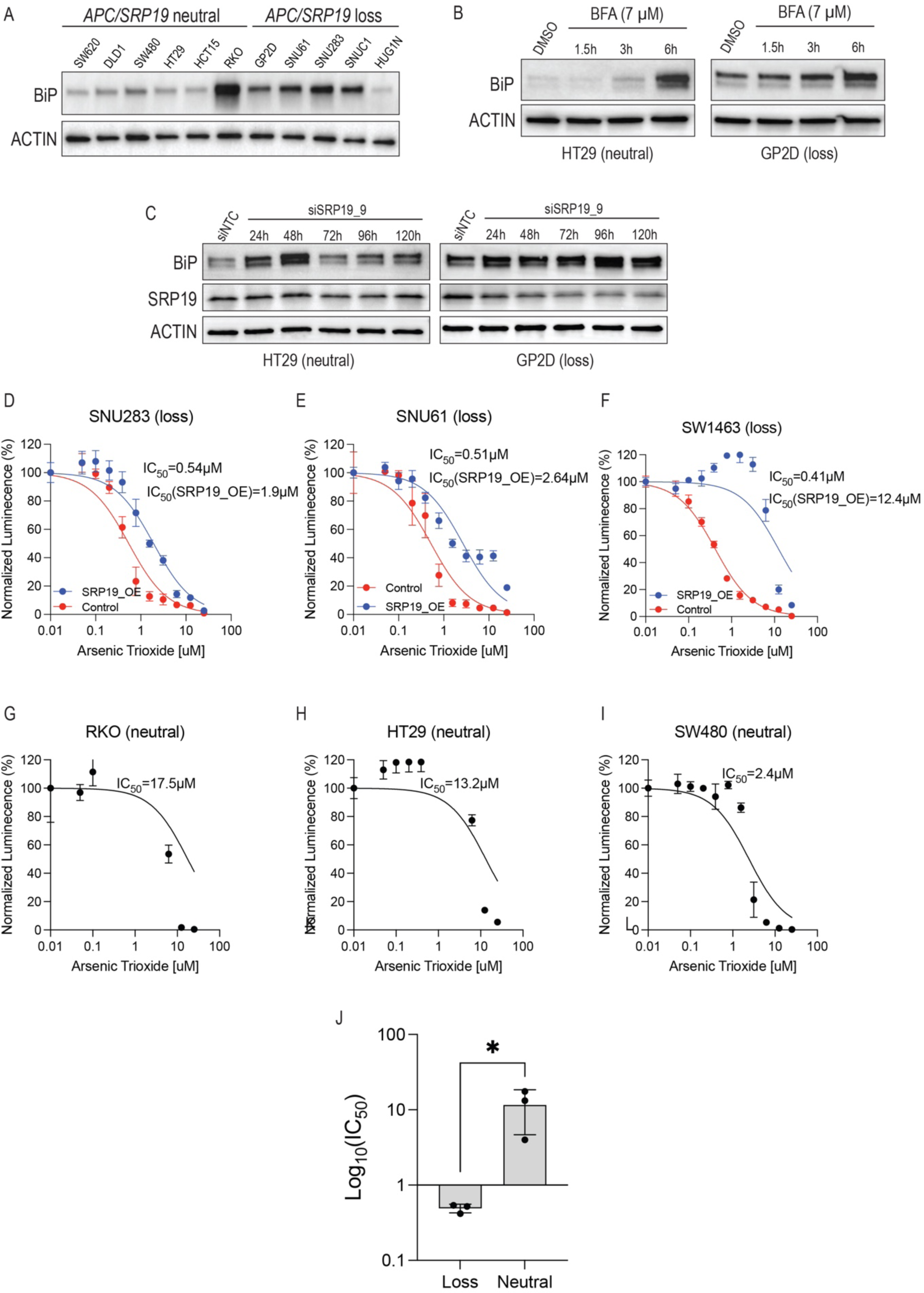
Cells with *APC/SRP19* loss have high basal levels of ER stress and are sensitive to Arsenic Trioxide. (A) BiP levels in cells with neutral or loss of *APC/SRP19* copy number. (B) BiP levels in HT29 (*APC/SRP19* neutral copy number) or GP2D (*APC/SRP19* copy number loss) following treatment with 7µM of BFA for the indicated time. (C) BiP levels in HT29 (*APC/SRP19* neutral copy number) or GP2D (*APC/SRP19* copy number loss) following transfection with SRP19 targeting siRNA. (D-F) Cell proliferation measured using cell titre glow 5 days post treatment with Arsenic Trioxide in *APC/SRP19* loss cells with or without *SRP19* overexpression. (G-I) Cell proliferation measured using cell titre glow 5 days post treatment with Arsenic Trioxide in *APC/SRP19* neutral cells. (J) Compression of Arsenic Trioxide IC50 in cells with neutral or loss of *APC/SRP19* copy number. p.Value calculated using a two-tailed unpaired T.test.

Molecules that induce ER stress are found in naturally occurring toxins and have potent effects on cell proliferation. Based on our observations we hypothesised that some of these toxins will have a more potent effect in *APC/SRP19* loss cells. Exotoxin A and Eeyarestatin I are toxins that interact with SEC61 and inhibit its activity(31). Using a panel of cell-lines we found that Exotoxin A (Supplementary Fig. 5A-C) and Eeyarestatin I (Supplementary Fig. 5D-F) inhibited proliferation in all cell lines regardless of *APC/SRP19* copy number alterations. BFA induces ER stress by inhibiting transport from the ER to the golgi. Treatment of cells with BFA had a toxic anti-proliferation effect that was not related to *APC/SRP19* copy number (Supplementary Fig. G-H).

Arsenic Trioxide is a clinically approved drug for relapsed leukemia that induces ER stress by causing oxidative stress and activating the ER stress response(32). Arsenic Trioxide treatment induced ER stress in *APC/SRP19* loss cells as indicated by high levels of the ER stress marker CHOP (Supplementary Fig. 5J). Following treatment with Arsenic Trioxide we measured proliferation in a panel of cells with neutral or loss of *APC/SRP19* copy number (Fig. 6D-J). At concentrations >1-5µM Arsenic Trioxide was toxic and had an anti-proliferation effect in all cell lines. At lower concentrations (0.4-0.5µM) we found that only cells harbouring loss of *APC/SRP19* copy number were sensitive to Arsenic Trioxide (Fig. 6D-F). The average IC_50_ for cells with neutral *APC/SRP19* copy number was 11.1µM (Fig. 6J) which is similar range to the Arsenic Trioxide IC_50_ reported in a large collection of cell lines(33). Cells harbouring *APC/SRP19* loss were highly sensitive to Arsenic Trioxide treatment and had an average IC_50_ of 0.48µM suggesting low dose Arsenic Trioxide as a potential treatment in these cells. To validate that the observed proliferation phenotype is due to the lower levels of *SRP19,* we used a rescue experiment. Following *SRP19* overexpression in SW1463, SNU283 or SNU61 cells were treated with Arsenic Trioxide and proliferation was measured after 5 days (Fig. 6D-F). *SRP19* overexpression shifted Arsenic Trioxide sensitivity to 1.9-12.4µM, like what we observed in cells with neutral *APC/SRP19* copy number (Fig. 6G-I). These results demonstrate that Arsenic Trioxide induction of ER stress in cells with *APC/SRP19* loss is mediated by low levels of *SRP19* and suggests the use of low dose Arsenic Trioxide as a treatment in these cancers.

## Discussion

*APC* is a TSG that is frequently mutated in cancer and is lost in ∼80% of colorectal cancers. Early studies have shown that loss of *APC* activity is one of the earliest events in colon cancer leading to benign polyp formation in humans and animal models(34). Furthermore, re-expression of *APC* abolished tumour growth in animal models even in advanced high-grade carcinomas(7) demonstrating that loss of *APC* is required for tumour maintenances. Yet, no current therapy is available for direct targeting of a TSG that is lost in cancer. Cancer sequencing studies(3) identified two main mechanisms for loss of *APC* activity, mutations in the protein coding region and heterozygous deletions. In many cases complete loss of *APC* is achieved by mutating one allele and deleting the other. In the current work we found that *SRP19*, a cell essential gene that is located 15kb away from *APC*, is co-deleted in cancers with heterozygous *APC* loss. The collateral loss of cell essential genes driven by proximity to a TSG may result in vulnerabilities that could be exploited for specific targeting of cancer.

Loss of function genetic screens are highly useful in identifying new targets for drug development(19,35). Recent efforts to map the landscape of genetic dependencies in cancer have focused on CRISPR knockout strategies since they have lower noise and less off target effects then RNAi(9,16,36). We show here that although RNAi has more off-target effects then CRISPR, partial gene dependencies could be identified only with RNAi and not with CRISPR based approaches. Specifically, CRISPR knockouts eliminate the target gene as opposed to RNAi which only reduce the levels of the target gene. When looking at cell essential genes RNAi provides a unique opportunity to find cancer targets that are dose dependent. Recent studies showed that RNAi but not CRISPR is uniquely capable of identifying gene-gene interactions in large scale loss of function proliferation screens for core cell essential genes(18).

Previous studies have identified genes targets that are deleted due their proximity to a TSG(8,11). In all these cases heterozygous deletion of the deleted gene did not lead to a dramatic difference in protein activity. For example, *PSMC2* is lost in many cancers and partial *PSMC2* knockdown is vulnerability in these cancers(8). However, proteosome activity was not affected in cells with loss of *PSMC2* and proteosome inhibitors were not effective at differential targeting of cells with *PSMC2* loss. In the current study we show that *SRP19* is the rate limiting factor for formation of the SRP complex and that loss of *SRP19* leads to loss of other components of the SRP complex and to lower activity of this complex. Due to this effect on SRP complex activity drugs that target the activity of this complex are effective in treatment of cells harbouring these *APC/SRP19* loss. Our study suggests that identifying genes that are rate limiting for formation of an active complex is more efficient in finding targetable genes. Different than other collateral loss genes *SRP19* unique location and proximity to *APC* ensure that *SRP19* is almost always lost together with *APC*. Our study demonstrates that the distance of a collateral loss gene from a TSG is important for identifying similar targets for other TSGs.

The SRP complex is a highly conserved protein-RNA structure that has evolved to regulate translocation of proteins to the ER(14). In the current study we found that one of these proteins, *SRP19*, is deleted due to its proximity to the TSG *APC*. *APC/SRP19* deletion leads to lower *SRP19* mRNA and protein and, as a consequence, these cells are highly sensitive to partial inhibition of *SRP19* expression. In addition, we found that *SRP19* levels regulate the stability of other SRP complex proteins. Lower levels of SRP19 protein leads to lower SRP54 protein stability and consequently lower activity of the SRP complex and protein secretion. Our results suggest that binding of SRP19 to 7SL complex is required for enabling binding of SRP54 and SRP68 and formation of an active SRP complex. Decreased complex stability due to low levels of a protein is a unique mechanism that has not been observed for other genes such as *PSMC2(8)* or *SF3B1*(12). In the case of *PSMC2* heterozygous deletion depletes a pool of *PSMC2* protein which serves as a reservoir. Here we showed that different to *PSMC2*, *SRP19* loss destabilises other components of the complex as well as the activity of the SRP complex. Our results suggest that genes that are part of multi-subunit complexes and are rate limiting for complex formation may be more potent targets.

In addition to proteins that are part of the S-domain *SRP9* which is part of the SRP-complex Alu domain was also a vulnerability in *APC/SRP19* loss cancers. Although we currently do not completely understand *SRP9* sensitivity it is possible that *SRP9* regulate other functions that are not related to the SRP complex and that lower SRP complex activity leads to depletion of the *SRP9* pool. Interestingly SRP9 has been shown to interact with Alu sequences that are not related to the SRP complex and may have additional roles(37).

We found that partial suppression of ribosomal genes was negatively associated with *SRP19* dependency. The SRP complex binds directly to the ribosome during translation and mediates the translocation of the emerging protein to the ER(25). These findings suggest that cancer cells with heterozygous loss of *APC/SRP19* are less dependent on partial inhibition of the ribosome. These observations could have important implications to the use of ribosome inhibitors for cancer treatment(38).

We found that low levels of SRP complex activity led to higher basal levels of ER stress measured by high basal levels of the protein chaperon BiP in *APC/SRP19* loss cells. These observations suggest the use of molecules that induce ER stress to treat these cancers. Exotoxin A, Eeyarestatin I and BFA inhibited proliferation in all cell lines and did not show any effect that was specific for *APC/SRP19* loss cells. It is likely that these potent toxins do not show differential sensitivity since they inhibit additional cellular pathways and signalling networks. We found that low concentrations of Arsenic Trioxide inhibited proliferation only in *APC/SRP19* loss cells and was rescued by overexpression of *SRP19*. Arsenic Trioxide is tolerated in humans and is an approved therapy for relapsed acute promyelocytic leukemia(39). Our results suggest that low dose Arsenic Trioxide is an effective therapy in these cancers. Although currently no drugs that directly target SRP19 or SRP54 or SRP68 are available future development of such molecules may have a better window for treating cancers with loss of *APC/SRP19*.

In summary, we demonstrate that *SRP19* mRNA and protein are lost in cancers that harbour *APC* loss. We show that *SPR19* loss destabilises the SRP complex and leads to lower activity of the SRP complex and decreased rate of protein secretion. Furthermore, we find that molecules that induce ER stress are highly effective in inhibiting growth of these cancers.

## Data Availability

All raw and processed data are available in the figures, supplementary figures, and supplementary tables of this manuscript. All plasmids generated in this study will be made available through Addgene.

## Methods

### Cell line and media

Cell-line identity was verified by STR at the Australian Genomics Research Facility (AGRF). Cancer cell lines were obtained from ATCC (SW620, DLD1, HCT15, SW480, HT29, RKO, GP2D, SNU283, SNU61, SW1463, SNUC1, HUG1N, SW1417). HEK293FT cells were Thermo-Fisher (Cat num#R70007). Cell lines were maintained in a 37°C incubator. All cell lines were propagated in media containing 10% FBS, 1% penicillin and Streptomycin and 1% L-glutamine in SW620, DLD1, HCT15, SW480, HT29, RKO and GP2D were cultured in DMEM (Thermo-Fisher Scientific#11965092), SW1463 were cultured in DMEM with 1mM Na-Pyruvate (Thermo-Fisher Scientific#11360070), SNUC1, HUG1N, SW1417 were cultured in RPMI-1640 medium (Thermo-Fisher Scientific#11875093), SNU283, SNU61 were cultured in RPMI-1640 with 25mM HEPES (Thermo-Fisher Scientific#15630080).

### RNA isolation and quantitative real-time PCR

Total RNA was isolated using TRIzol (Sigma-Aldrich) from mouse tumours and cell lines, according to the manufacturer’s instructions. One microgram of total RNA was reverse transcribed to cDNA using the Maxima Reverse Transcriptase (Thermo-Fisher#EP0742) in 20μl reaction with random hexamers and dNTP. Expression levels of genes were quantified using qRT-PCR using the QuantStudio™5 Real-Time PCR system (Thermo-Fisher). qRT-PCR was performed using the indicated qPCR primers (Supplementary Table S3). ACTIN or GAPDH primers were used for normalisation. The qRT-PCR conditions were denaturation at 95°C for 5min followed by 40 cycles of amplification at 95°C for 15s and 60°C for 20s. The comparative cycle threshold (delta Ct) method was used to analyse the gene expression levels.

### siRNA transfection

siRNAs (Supplementary Table S3) were obtained from IDT which and designed to target SRP proteins including *SRP19*, *SRP14*, *SRP9*, *SRP54*, *SRP68* and *SRP72*. Lipofectamine RNAiMAX (Invitrogen#13778150) was used for siRNA transfection according to the manufacturer’s instructions.

### Crystal violet proliferation assay

Following siRNA transfection or drug treatment, cells were allowed to propagate for the indicated time. The media was removed, and cells were washed twice in PBS. 10% of formalin in PBS was added and incubated for 20min at room temperature. Formalin was removed, and 0.5% (w/v) of crystal violet (Sigma#C0775-25G) was added and incubated for 20min at room temperature. Crystal violet was removed, and plates were thoroughly washed with PBS. For quantification, 10% of acetic acid was added to each well and incubated at room temperature for 3min. The extracted solution was added to a 96-well plate and quantified by measuring the OD at 590nm.

### Protein quantification using Western blot

Cells/tissue were harvested, washed in PBS, and resuspended in RIPA buffer (CST-9806) containing proteinase inhibitors (Roche) and quantified using the Pierce BCA Protein Assay Kit (Thermo-Fisher#23225). Protein lysates diluted in 4 X Laemmli Sample Buffer (Bio-Rad#161–0747) were loaded onto Bio-Rad 4–20% precast gels. Western blot was done using BioRad pre-casted gels and a trans-turbo transfer machine (BioRad). Following electrophoresis, proteins were transferred to a pre-activated PVDF membrane using the Trans-Blot®Turbo™ Transfer System and visualized using ECL (Bio-Rad Chemidoc). The following are the antibodies used in this study: SRP72 (Sigma#034621), SRP68 (Invitrogen#100080), SRP54 (BD Bioscience #610940), SRP19 (Invitrogen #93197), ACTIN (Santa Cruz#A2017), GAPDH (Sana Cruz # SC32233), BiP (Cell Signaling#3183).

### SEAP enzymatic assay

Cells were transduced with a lentivirus SEAP expression. Following puromycin selection (3 DPI), cells were seeded on 24-well plate for siRNA transfection or drug treatment. For siRNA treatment, cell culture media was collected after 72h post transfection. For drug treatment, cell culture media was collected after 6h. Secreted SEAP were detected using the Great EscAPe™ SEAP Chemiluminescence Kit 2.0 (Clontech#631738) following the manufacturer’s instructions. Live cell counts were measured using an Invitrogen Countess 3 Automated Cell Counter, and the secreted SEAP level was normalized to the corresponding cell counts.

### Mouse xenografts

All animal studies were approved by the Monash University Animal Ethics Committee (AEC-approval number 2020-24197-49078). RKO and GP2D cells were infected with lentiviruses containing the inducible Tet-on vector expressing shRNA targeting SRP19 when adding doxycycline. Following puromycin selection (3 DPI), 1e6 cells/site and 3 sites/mouse were subcutaneously injected into 5-week-old female NSG mice under isoflurane anaesthesia. In the experimental groups, mice were fed with doxycycline food (600mg/kg). Tumour growth was continuously monitored for 4 weeks. Tumours were measured using digital vernier calliper every 48h, beginning 3 days after injection, and tumour volume was calculated using the formula length (mm)×width (mm)×height (mm) and expressed in mm^3^.

## Acknowledgments

This work was supported by an NHMRC grant (grant number: 1140808) to J.R. and a Tour de-Cure grant to J.R., E.S and M.C. (grant number: RSP-225-18/19). J.R is supported by a Victoria Cancer Agency fellowship (grant number: MCRF20035). We thank the Functional Genomics Platform and the Genomics and Bioinformatics Platform at Monash University for help with data analysis. We thank Prof. Renea Taylor (Monash University) for providing mice. We thank Prof. Lee Wong (Monash University) for providing Arsenic Trioxide.

## Supplementary Figures

**Supplementary Figure 1:**
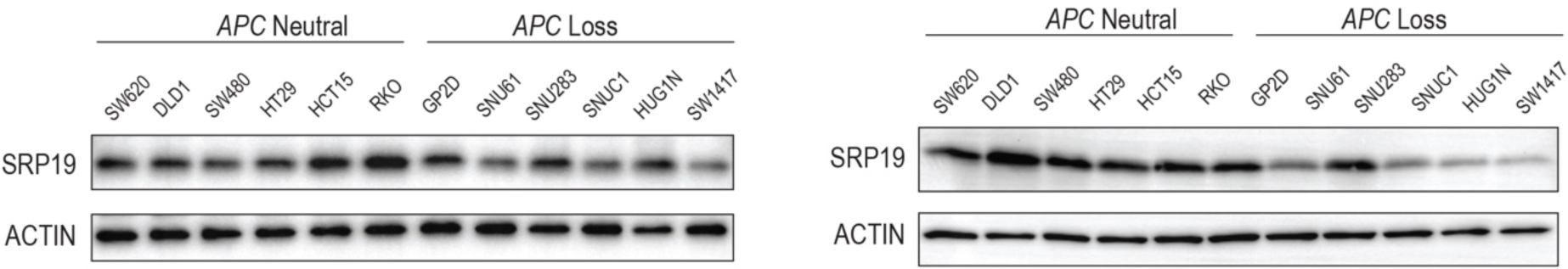
*APC* loss leads to loss of *SRP19* DNA, mRNA and protein. Biological replicates quantifying SRP19 protein levels.

**Supplementary Figure 2:**
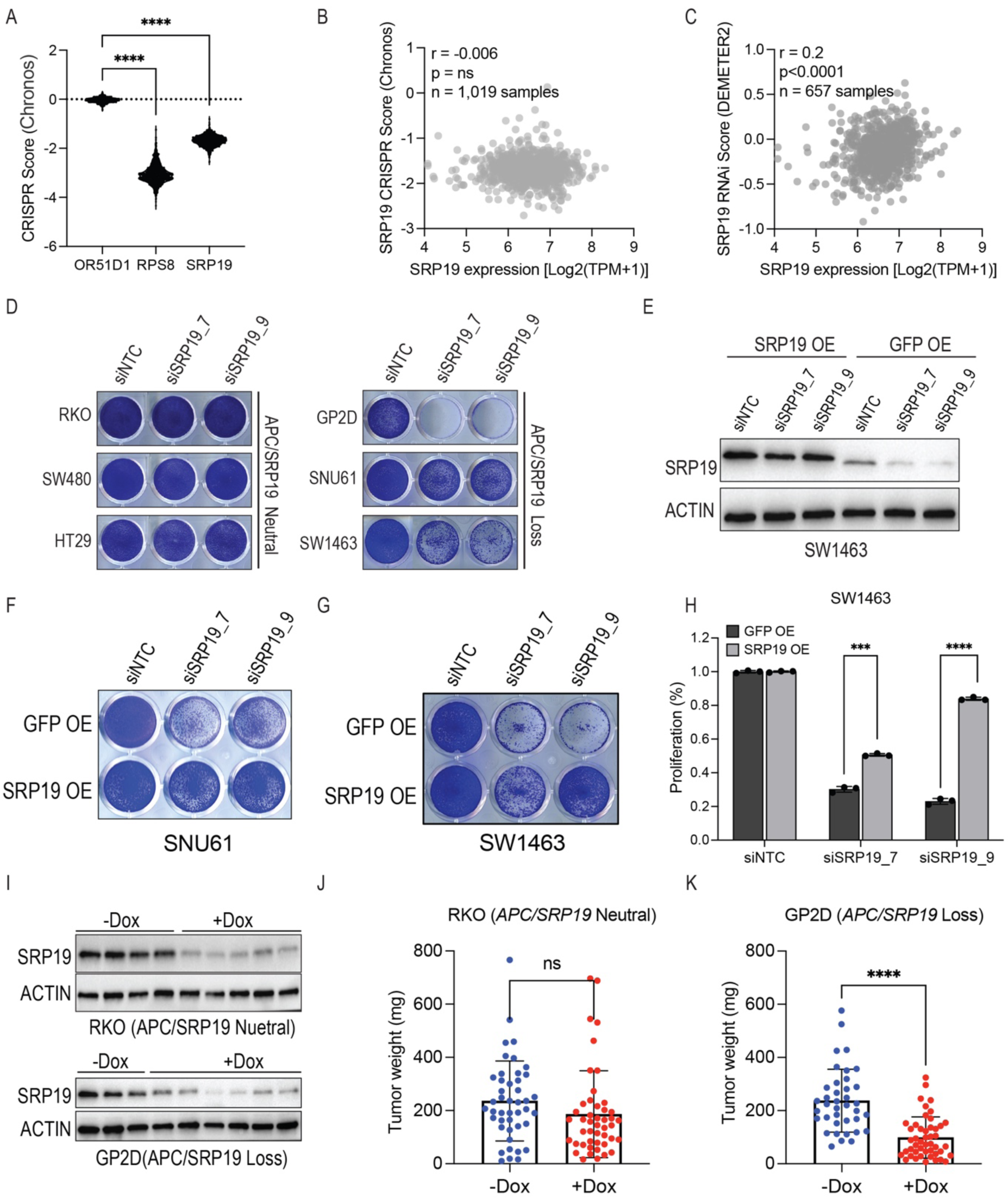
Partial inhibition of *SRP19* is a vulnerability in cells with *APC/SRP19* loss. (A) Proliferation following CRISPR mediated knockout of *SRP19* from DepMap. *OR51D1* is an olfactory receptor and used as a negative control. *RPS8* is a core essential ribosomal gene and used as a positive control. p.Value calculated using two tailed unpaired T.test. (B) Correlation between *SRP19* mRNA expression and proliferation using CRISPR knockout, data from DepMap. (C) Correlation between *SRP19* mRNA expression and proliferation using shRNA, data from DepMap. (D) Crystal violet images of *APC/SRP19* neutral or loss cells 7 days post transfection with SRP19 targeting siRNAs. SNUC1 and HUG1N are suspension cell lines and proliferation, in these cell lines, was measured using a cell counter. (E) SRP19 protein levels in SW1463 cells overexpressing GFP (control) or *SRP19*, 3 days post transfection with *SRP19* targeting siRNAs. (F) Crystal violet images of SNU61 cells overexpressing *SRP19* or GFP 7 days post transfection with *SRP19* targeting siRNAs. (G) Crystal violet image of SW1463 cells overexpressing GFP or *SRP19* 7 days post transfection with *SRP19* targeting siRNAs. (H) Quantification of crystal violet images in (G). (I) SRP19 protein levels in tumours extracted from mice treated with or without DOX. (J) Tumour weight of tumours from mice injected with RKO cells. p.Value calculated using two tailed unpaired T.test. (K) Tumour weight of tumours from mice injected with GP2D cells. p.Value calculated using two tailed unpaired T.test.

**Supplementary Figure 3:**
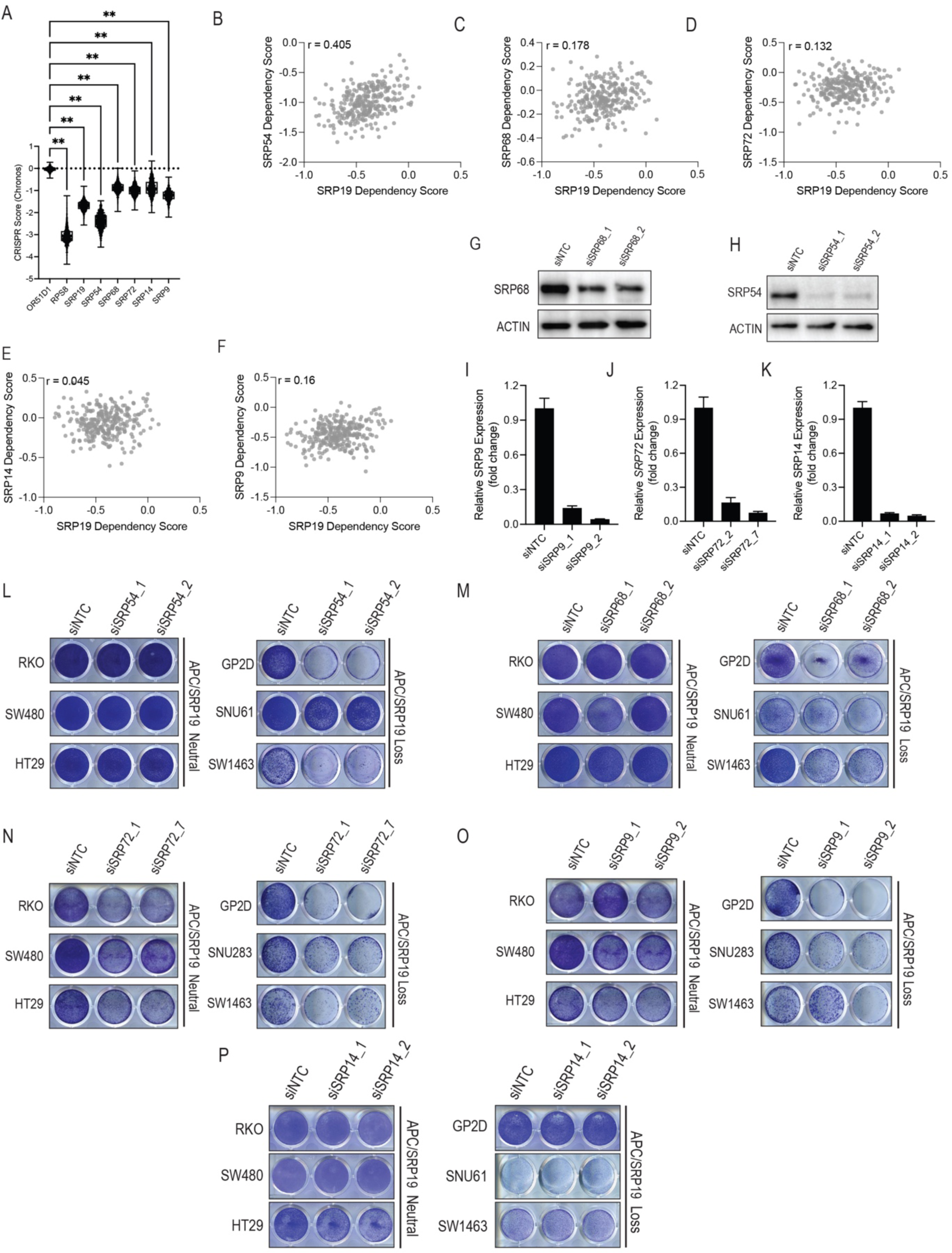
*SRP68* and *SRP54* are also vulnerabilities in *APC/SRP19* loss cancers. Comparison of *SRP19* RNAi dependency scores across 648 cancer cell lines (DepMap) with (A) *SRP54*. (B) *SRP68*. (C) *SRP72*. (D) *SRP14*. (E) *SRP9*. (F) Proliferation following CRISPR mediated knockout of SRP complex proteins in 970 cell lines (DepMap). *OR51D1* is a negative control and *RPS8* is a ribosomal core essential gene and used as a positive control. pValue calculated using two tailed unpaired T.test. (G) Protein or mRNA levels following siRNA mediated suppression of SRP complex proteins: (G) SRP68. (H) SRP54. (I) *SRP9*. (J) *SRP72*. (K) *SRP14*. Crystal violet images (SNUC1 and HUG1N are suspension lines and proliferation was monitored by counting) of *APC/SRP19* loss or neutral cells 7 days post transfection with siRNAs targeting: (L) *SRP54.* (M) *SRP68.* (N) *SRP72.* (O) *SRP9.* (P) *SRP14*.

**Supplementary Figure 4:**
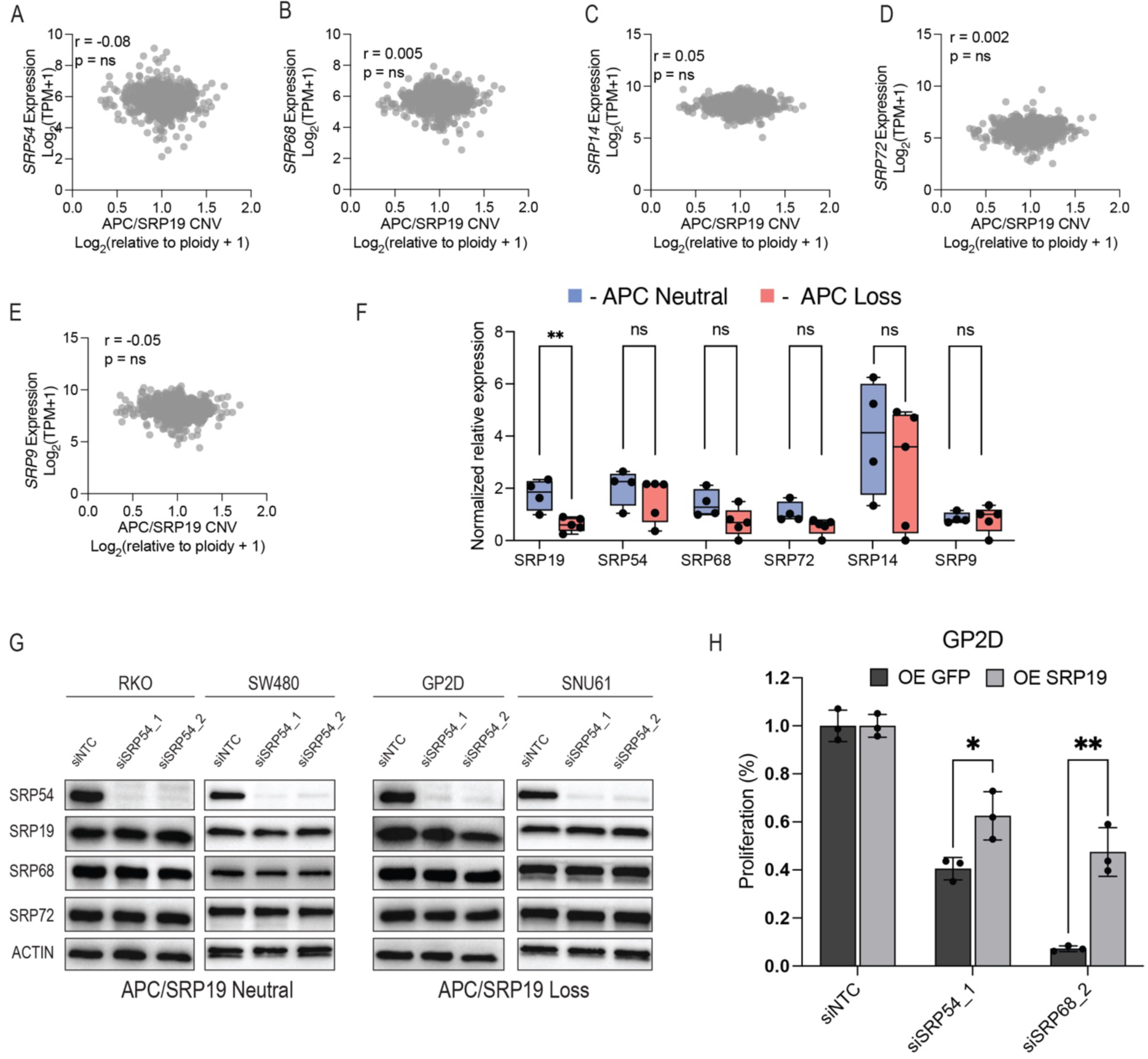
Loss of *APC/SRP19* destabilises components of the SRP complex. Pearson correlation across 1,440 CCLE cell lines of *APC/SRP19* copy number and mRNA expression of (A) *SRP54* (B) *SRP68* (C) *SRP14* (D) *SRP72* (E) *SRP9* (F) Expression of the indicated genes was measured using qRT-PCR in cells with neutral or loss of *APC/SRP19* copy number. pValue calculated using two tailed unpaired T.test. (G) Protein levels of SRP complex components following transfection of cell lines with neutral or loss of *APC/SRP19* copy number with *SRP54* targeting siRNAs. (H) Proliferation of GP2D cells following overexpression of SRP19 and transfection of *SRP54* targeting siRNAs. pValue calculated using two tailed unpaired T.test.

**Supplementary Figure 5:**
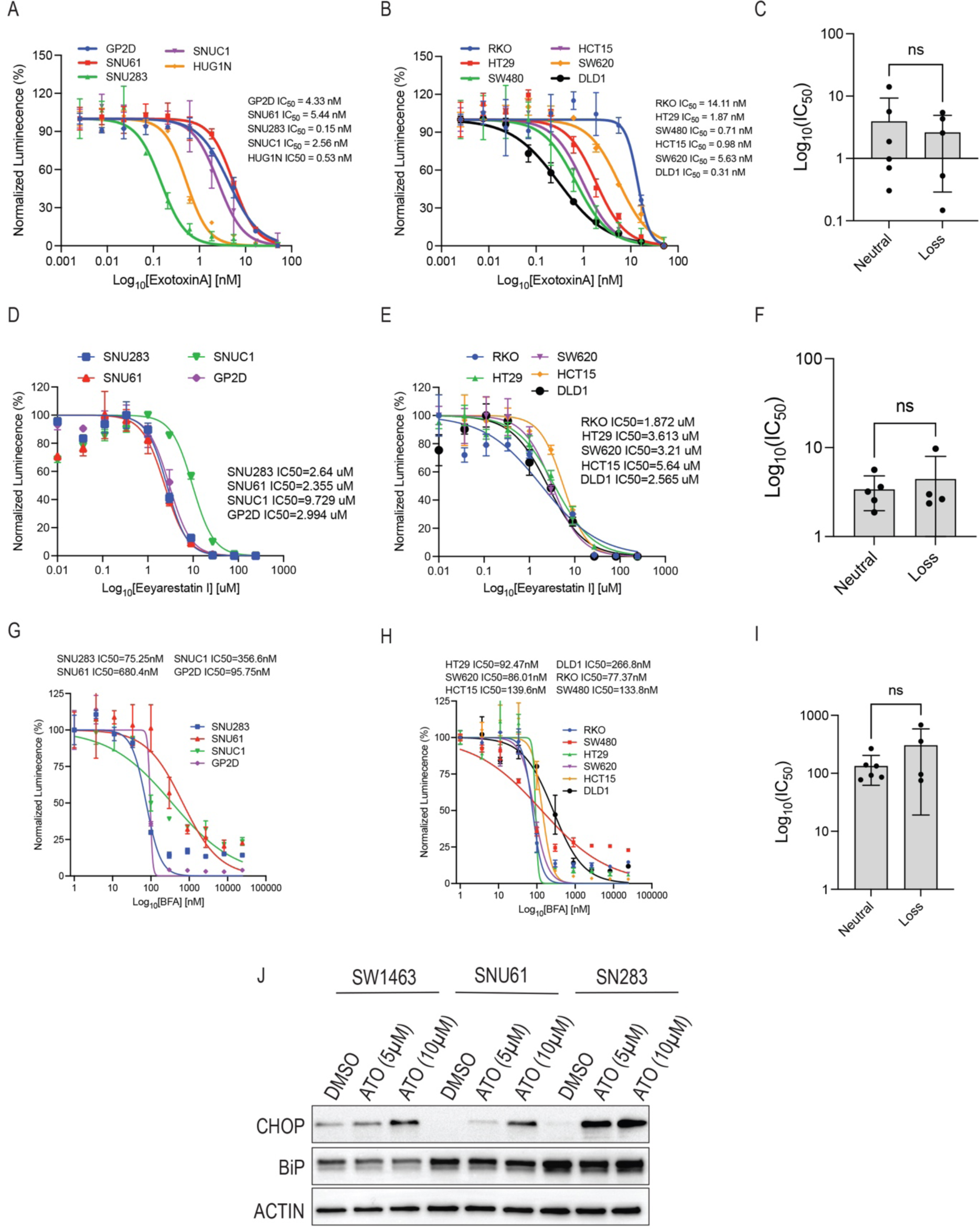
Effect of molecules that inhibit the protein secretion on proliferation of *APC/SRP19* loss cells. (A) Dose inhibition curve measured using CTG following treatment of cells with *APC/SRP19* copy number loss with Exotoxin A. (B) Dose inhibition curve measured using CTG following treatment of cells with neutral *APC/SRP19* copy number with Exotoxin A. (C) Comparison of Exotoxin A IC50 in cells with neutral or loss of *APC/SRP19* copy number. pValue calculated using two tailed unpaired T.test. (D) Dose inhibition curve measured using CTG following treatment of cells with *APC/SRP19* copy number loss with Eeyarestatin I. (E) Dose inhibition curve measured using CTG following treatment of cells with neutral *APC/SRP19* copy number with Eeyarestatin I. (F) Comparison of Eeyarestatin I IC50 in cells with neutral or loss of *APC/SRP19* copy number. pValue calculated using two tailed unpaired T.test. (G) Dose inhibition curve measured using CTG following treatment of cells with *APC/SRP19* copy number loss with BFA. (H) Dose inhibition curve measured using CTG following treatment of cells with neutral *APC/SRP19* copy number with BFA. (I) Comparison of BFA IC50 in cells with neutral or loss of *APC/SRP19* copy number. pValue calculated using two tailed unpaired T.test. (J) Levels of the ER stress markers BiP and CHOP 24h after treatment with Arsenic Trioxide (ATO).

## Supplementary Table Legends

**Supplementary Table S1: Pearson correlation values of APC CNV with gene expression changes across the CCLE data set (1,440 cell lines).**

**Supplementary Table S2: Pearson correlation values of SRP19 dependency (RNAi) and gene dependencies across the DepMap data set (657 cell lines).**

**Supplementary Table S3: Primers and RNAi sequences used in this study.**

## References

1. Clevers H, Nusse R. Wnt/beta-catenin signaling and disease. Cell 2012;149(6):1192–205 doi 10.1016/j.cell.2012.05.012.

2. Rosenbluh J, Wang X, Hahn WC. Genomic insights into WNT/beta-catenin signaling. Trends Pharmacol Sci 2014;35(2):103–9 doi 10.1016/j.tips.2013.11.007.

3. Cancer Genome Atlas N. Comprehensive molecular characterization of human colon and rectal cancer. Nature 2012;487(7407):330-7 doi 10.1038/nature11252.

4. Fang DC, Luo YH, Yang SM, Li XA, Ling XL, Fang L. Mutation analysis of APC gene in gastric cancer with microsatellite instability. World J Gastroenterol 2002;8(5):787–91 doi 10.3748/wjg.v8.i5.787.

5. Furuuchi K, Tada M, Yamada H, Kataoka A, Furuuchi N, Hamada J, et al. Somatic mutations of the APC gene in primary breast cancers. Am J Pathol 2000;156(6):1997–2005 doi 10.1016/s0002-9440(10)65072-9.

6. Vasaikar S, Huang C, Wang X, Petyuk VA, Savage SR, Wen B, et al. Proteogenomic Analysis of Human Colon Cancer Reveals New Therapeutic Opportunities. Cell 2019;177(4):1035–49 e19 doi 10.1016/j.cell.2019.03.030.

7. Dow LE, O’Rourke KP, Simon J, Tschaharganeh DF, van Es JH, Clevers H, et al. Apc Restoration Promotes Cellular Differentiation and Reestablishes Crypt Homeostasis in Colorectal Cancer. Cell 2015;161(7):1539–52 doi 10.1016/j.cell.2015.05.033.

8. Nijhawan D, Zack TI, Ren Y, Strickland MR, Lamothe R, Schumacher SE, et al. Cancer vulnerabilities unveiled by genomic loss. Cell 2012;150(4):842–54 doi 10.1016/j.cell.2012.07.023.

9. Rosenbluh J, Mercer J, Shrestha Y, Oliver R, Tamayo P, Doench JG, et al. Genetic and Proteomic Interrogation of Lower Confidence Candidate Genes Reveals Signaling Networks in beta-Catenin-Active Cancers. Cell Syst 2016;3(3):302–16 e4 doi 10.1016/j.cels.2016.09.001.

10. Liu Y, Zhang X, Han C, Wan G, Huang X, Ivan C, et al. TP53 loss creates therapeutic vulnerability in colorectal cancer. Nature 2015;520(7549):697-701 doi 10.1038/nature14418.

11. Muller FL, Colla S, Aquilanti E, Manzo VE, Genovese G, Lee J, et al. Passenger deletions generate therapeutic vulnerabilities in cancer. Nature 2012;488(7411):337-42 doi 10.1038/nature11331.

12. Paolella BR, Gibson WJ, Urbanski LM, Alberta JA, Zack TI, Bandopadhayay P, et al. Copy-number and gene dependency analysis reveals partial copy loss of wild-type SF3B1 as a novel cancer vulnerability. Elife 2017;6 doi 10.7554/eLife.23268.

13. Kronke J, Fink EC, Hollenbach PW, MacBeth KJ, Hurst SN, Udeshi ND, et al. Lenalidomide induces ubiquitination and degradation of CK1alpha in del(5q) MDS. Nature 2015;523(7559):183-8 doi 10.1038/nature14610.

14. Wild K, Becker MMM, Kempf G, Sinning I. Structure, dynamics and interactions of large SRP variants. Biol Chem 2019;401(1):63–80 doi 10.1515/hsz-2019-0282.

15. Barretina J, Caponigro G, Stransky N, Venkatesan K, Margolin AA, Kim S, et al. The Cancer Cell Line Encyclopedia enables predictive modelling of anticancer drug sensitivity. Nature 2012;483(7391):603-7 doi 10.1038/nature11003.

16. Ghandi M, Huang FW, Jane-Valbuena J, Kryukov GV, Lo CC, McDonald ER, 3rd, et al. Next-generation characterization of the Cancer Cell Line Encyclopedia. Nature 2019;569(7757):503-8 doi 10.1038/s41586-019-1186-3.

17. Cerami E, Gao J, Dogrusoz U, Gross BE, Sumer SO, Aksoy BA, et al. The cBio cancer genomics portal: an open platform for exploring multidimensional cancer genomics data. Cancer Discov 2012;2(5):401–4 doi 10.1158/2159-8290.CD-12-0095.

18. Krill-Burger JM, Dempster JM, Borah AA, Paolella BR, Root DE, Golub TR, et al. Partial gene suppression improves identification of cancer vulnerabilities when CRISPR-Cas9 knockout is pan-lethal. Genome Biol 2023;24(1):192 doi 10.1186/s13059-023-03020-w.

19. Tsherniak A, Vazquez F, Montgomery PG, Weir BA, Kryukov G, Cowley GS, et al. Defining a Cancer Dependency Map. Cell 2017;170(3):564–76 e16 doi 10.1016/j.cell.2017.06.010.

20. Wiederschain D, Wee S, Chen L, Loo A, Yang G, Huang A, et al. Single-vector inducible lentiviral RNAi system for oncology target validation. Cell Cycle 2009;8(3):498–504 doi 10.4161/cc.8.3.7701.

21. Pan J, Meyers RM, Michel BC, Mashtalir N, Sizemore AE, Wells JN, et al. Interrogation of Mammalian Protein Complex Structure, Function, and Membership Using Genome-Scale Fitness Screens. Cell Syst 2018;6(5):555–68 e7 doi 10.1016/j.cels.2018.04.011.

22. Subramanian A, Tamayo P, Mootha VK, Mukherjee S, Ebert BL, Gillette MA, et al. Gene set enrichment analysis: a knowledge-based approach for interpreting genome-wide expression profiles. Proc Natl Acad Sci U S A 2005;102(43):15545–50 doi 10.1073/pnas.0506580102.

23. Grotwinkel JT, Wild K, Segnitz B, Sinning I. SRP RNA remodeling by SRP68 explains its role in protein translocation. Science 2014;344(6179):101-4 doi 10.1126/science.1249094.

24. Becker MM, Lapouge K, Segnitz B, Wild K, Sinning I. Structures of human SRP72 complexes provide insights into SRP RNA remodeling and ribosome interaction. Nucleic Acids Res 2017;45(1):470–81 doi 10.1093/nar/gkw1124.

25. Halic M, Blau M, Becker T, Mielke T, Pool MR, Wild K, et al. Following the signal sequence from ribosomal tunnel exit to signal recognition particle. Nature 2006;444(7118):507-11 doi 10.1038/nature05326.

26. Kuglstatter A, Oubridge C, Nagai K. Induced structural changes of 7SL RNA during the assembly of human signal recognition particle. Nat Struct Biol 2002;9(10):740–4 doi 10.1038/nsb843.

27. Lakkaraju AK, Luyet PP, Parone P, Falguieres T, Strub K. Inefficient targeting to the endoplasmic reticulum by the signal recognition particle elicits selective defects in post-ER membrane trafficking. Exp Cell Res 2007;313(4):834–47 doi 10.1016/j.yexcr.2006.12.003.

28. Lowe ME. Site-specific mutations in the COOH-terminus of placental alkaline phosphatase: a single amino acid change converts a phosphatidylinositol-glycan-anchored protein to a secreted protein. J Cell Biol 1992;116(3):799–807 doi 10.1083/jcb.116.3.799.

29. Chardin P, McCormick F. Brefeldin A: the advantage of being uncompetitive. Cell 1999;97(2):153–5 doi 10.1016/s0092-8674(00)80724-2.

30. Bertolotti A, Zhang Y, Hendershot LM, Harding HP, Ron D. Dynamic interaction of BiP and ER stress transducers in the unfolded-protein response. Nat Cell Biol 2000;2(6):326–32 doi 10.1038/35014014.

31. Linxweiler M, Schick B, Zimmermann R. Let’s talk about Secs: Sec61, Sec62 and Sec63 in signal transduction, oncology and personalized medicine. Signal Transduct Target Ther 2017;2:17002 doi 10.1038/sigtrans.2017.2.

32. Wadgaonkar P, Chen F. Connections between endoplasmic reticulum stress-associated unfolded protein response, mitochondria, and autophagy in arsenic-induced carcinogenesis. Semin Cancer Biol 2021;76:258–66 doi 10.1016/j.semcancer.2021.04.004.

33. Sertel S, Tome M, Briehl MM, Bauer J, Hock K, Plinkert PK, et al. Factors determining sensitivity and resistance of tumor cells to arsenic trioxide. PLoS One 2012;7(5):e35584 doi 10.1371/journal.pone.0035584.

34. Powell SM, Zilz N, Beazer-Barclay Y, Bryan TM, Hamilton SR, Thibodeau SN, et al. APC mutations occur early during colorectal tumorigenesis. Nature 1992;359(6392):235-7 doi 10.1038/359235a0.

35. Howard TP, Vazquez F, Tsherniak A, Hong AL, Rinne M, Aguirre AJ, et al. Functional Genomic Characterization of Cancer Genomes. Cold Spring Harb Symp Quant Biol 2016;81:237–46 doi 10.1101/sqb.2016.81.031070.

36. Hahn WC, Bader JS, Braun TP, Califano A, Clemons PA, Druker BJ, et al. An expanded universe of cancer targets. Cell 2021;184(5):1142–55 doi 10.1016/j.cell.2021.02.020.

37. Bovia F, Fornallaz M, Leffers H, Strub K. The SRP9/14 subunit of the signal recognition particle (SRP) is present in more than 20-fold excess over SRP in primate cells and exists primarily free but also in complex with small cytoplasmic Alu RNAs. Mol Biol Cell 1995;6(4):471–84 doi 10.1091/mbc.6.4.471.

38. Kang J, Brajanovski N, Chan KT, Xuan J, Pearson RB, Sanij E. Ribosomal proteins and human diseases: molecular mechanisms and targeted therapy. Signal Transduct Target Ther 2021;6(1):323 doi 10.1038/s41392-021-00728-8.

39. Alimoghaddam K. A review of arsenic trioxide and acute promyelocytic leukemia. Int J Hematol Oncol Stem Cell Res 2014;8(3):44–54.

